# PiE: An open-source pipeline for home cage behavioral analysis

**DOI:** 10.1101/2023.05.05.539097

**Authors:** Jessie Benedict, Robert H Cudmore

**Author notes:** **Corresponding author:** Robert H Cudmore, PhD, Assistant Professor, Department of Physiology & Membrane Biology, School of Medicine, University of California - Davis, Address: Tupper Hall, Room 4136, One Shields Avenue, Davis, CA 95616.

## Abstract

Over the last two decades a growing number of neuroscience labs are conducting behavioral assays in rodents. The equipment used to collect this behavioral data must effectively limit environmental and experimenter disruptions, to avoid confounding behavior data. Proprietary behavior boxes are expensive, offer limited compatible sensors, and constrain analysis with closed-source hardware and software. Here, we introduce PiE, an open-source, end-to-end, user-configurable, scalable, and inexpensive behavior assay system. The PiE system includes the custom-built behavior box to hold a home cage, as well as software enabling continuous video recording and individual behavior box environmental control. To limit experimental disruptions, the PiE system allows the control and monitoring of all aspects of a behavioral experiment using a remote web browser, including real-time video feeds. To allow experiments to scale up, the PiE system provides a web interface where any number of boxes can be controlled, and video data easily synchronized to a remote location. For the scoring of behavior video data, the PiE system includes a standalone desktop application that streamlines the blinded manual scoring of large datasets with a focus on quality control and assay flexibility. The PiE system is ideal for all types of behavior assays in which video is recorded. Users are free to use individual components of this setup independently, or to use the entire pipeline from data collection to analysis. Alpha testers have included scientists without prior coding experience. An example pipeline is demonstrated with the PiE system enabling the user to record home cage maternal behavior assays, synchronize the resulting data, conduct blinded scoring, and import the data into R for data visualization and analysis.

## Introduction

Modern neuroscientific tools have enabled the manipulation of molecular and cellular function to then evaluate the effect on behavior. As the acquisition and analysis of awake and freely moving behaviors become more widespread, video capture becomes a critical mode of data collection in neuroscience.

The acquisition of this behavioral data introduces a number of challenges that must be overcome. It is now well established that environmental disruptions and experimenter presence can contaminate behavioral assays (Spruijt and DeVisser, 2006; Richardson, 2015; Krakauer *et al*., 2017; Voikar and Gaburro, 2020; Grieco *et al*., 2021). There is growing consensus that behavioral assays need to be standardized with respect to these disruptions to allow more meaningful comparisons between studies. To “place behavior on the same quantitative footing as other scientific fields” (Berman, 2018), behavior apparatuses should allow observations within a controlled and familiar home cage, while limiting the presence of other researchers in a shared vivarium, and to additionally limit the presence of an experimenter during behavioral data collection.

Many commercially available home cage behavioral systems are available, yet, these are often expensive, difficult to scale, and provide limited flexibility to incorporate the requirements for a particular behavioral experiment. As an alternative, a growing number of custom-built home cage monitoring systems have been developed (Goulding *et al*., 2008; Bains *et al*., 2018). One strategy is to retrofit a vivarium home cage with video monitoring (Salem *et al*., 2015; Singh *et al*., 2019). Alternative strategies to video recording of home cage activity monitoring have also been developed including implanted and external RFID chips (Bains *et al*., 2016; Redfern *et al*., 2017) and microwave sensors (Genewsky *et al*., 2017). Finally, there have been systems developed to monitor and record home cage learning paradigms (Balzani *et al*., 2018).

Once behavioral data is collected, it must be analyzed to extract the desired measurements and metrics required for a given assay. To achieve this, there are a growing number of video annotation software systems (See review: Luxem *et al*., 2022). Increasingly, this analysis software utilizes machine learning algorithms (See review: Brown and Bivort, 2018). An excellent example of machine learning applied to video recordings that can extract detailed pose information is DeepLabCut (Mathis *et al*., 2018; Nath *et al*., 2019; Lauer *et al*., 2022, Winters *et al*., 2022). A benefit of these machine learning algorithms is they can be run on any home cage video recordings.

Here we present PiE, an open-source and user-configurable home cage behavior pipeline from video capture to scored behavior for researchers with little to no programming experience. This system is inexpensive to build, will work equally well for just one home cage, and will scale easily to any number of home cages. A key advantage of this system is that the open-source software and accessible hardware affords the user flexibility to expand the behavior box function to meet the customized goals of their experiments. Using the provided browser-based software, the PiE system reduces environmental and experimenter disruptions by providing fully remote control and monitoring of home cage behaviors. Finally, the provided desktop GUI video annotation software, VideoAnnotate, streamlines manual and unbiased annotation of video recordings.

To test the PiE pipeline, we examined a complex social behavior by collecting video recordings and then analyzing maternal behavior of mouse dams from two inbred mouse lines and their pups over the course of weeks using 8 behavior boxes, recording simultaneously. Our PiE pipeline provides a robust end-to-end system for a range of behavioral analyses.

## Methods

### Animals

All animals were DBA/2J (Strain #:000671, RRID: IMSR_JAX:00067) or C57BL/6J (Strain #:000664, RRID: IMSR_JAX:000664) mice obtained from Charles River. Females (6-8 weeks old) used in behavior experiments were bred to males (8-10 weeks old) of their same strain. Female mice were group housed before and after breeding until approximately E17, when they were moved to single housing in a behavior box home cage prior to birth. All experiments were approved by the Animal Care and Use Committees of Johns Hopkins University. Animals were housed on a 12:12 light/dark cycle. Food and water were available *ad libitum*.

### Behavior Boxes

Behavior boxes were designed to enclose standard mouse cages (maximum of 5 mice per cage). The box was constructed with an aluminum frame (20 mm T-Slotted Rails, 80/20) and plastic walls (6 mm ABS Hair Panel, 80/20). Each box was equipped with a hinged door on the front, large enough to accommodate the cage and water bottle. Holes were drilled in the walls of the box to allow wires for sensors and actuators. Care was taken to make the boxes light tight and gaffer tape (25-50 mm black masking tape, Thor Labs) was used when necessary.

For our experiments, we included a number of sensors and actuators. Additional experimental components can easily be added as needed. We included (1) a ceiling mounted, downward facing, infrared (IR) video camera to allow both daytime and nighttime video recording, (2) white and IR LEDs to illuminate the behavior box (Note: 940 nm or shorter wavelength IR LEDs are preferred for optimal nighttime illumination), (3) a temperature and humidity sensor, and (4) a circulating fan for climate stability. The IR camera has a flex cable connecting it directly to the Raspberry Pi computer. The white and IR LEDS as well as the fan were powered via a quad-channel relay allowing them to be independently turned on and off by the Raspberry Pi computer. The tops of the home cages were replaced with a clear acrylic panel to provide an open field of view and improve the quality of the video recording. Mirrors were placed on either side of each mouse cage to allow the ceiling-mounted camera to capture side view video as well. These strategies greatly improved the over-all video quality and allowed the precise scoring of complex maternal behaviors.

We used boxes of approximately 24” W x 24” D x 24” H, but strongly recommend setting up the mouse cage, water bottle, and mirrors to confirm width and length dimensions. Likewise, we recommend setting up the ceiling mounted IR camera to determine the video field-of-view before deciding on the height of the box.

All components including suppliers, price, and additional notes are provided in Materials and Equipment (**Supplemental Table 1**).

### Raspberry Pi Computer

Each behavior box had a dedicated Raspberry Pi computer (Model 3B or newer, Raspberry Pi Foundation) to control the sensors and actuators within the box and to provide a web- and file-server to remotely control the experiment. Each Raspberry Pi computer included the computer itself, a 5V/2A AC/DC power adapter, a 16-32 GB micro SD card for the operating system, a 64-128 GB USB thumb drive to locally store recorded files including video, and an ethernet cable. Each computer also had a ribbon cable to connect the camera to the computer (2 m Flex Cable, Adafruit). One dedicated Raspberry Pi computer was attached to the back side of each behavior box.

Each Raspberry Pi computer used the Raspberry Pi OS operating system (version Buster). This OS is based on the Debian operating system which provides stable and efficient operation, making it well suited for the demands of long behavioral experiments and running on low-cost hardware such as the Raspberry Pi.

### Software Availability

All software is open-source and available on GitHub (https://github.com/cudmore/pie). Both technical and end user documentation is also provided including detailed software installation and troubleshooting for commonly encountered problems (https://cudmore.github.io/pie-doc/).

### PiE Server Software

We implemented custom software to control all aspects of a behavioral experiment. This software runs on a Raspberry Pi computer and once the hardware is configured can be installed with simple to use installation scripts. Implemented with a client/server architecture with a backend written in Python, a webserver (Flask), and a frontend web interface written in Javascript. We provide a point and click graphical user interface (GUI) that can run in any web-browser including on desktops, tablets, and smart-phones. All aspects of the software can be configured and controlled through a RESTful application-programming-interface (API) using Uniform Resource Locators (URLs) directly from a browser or with a programming language such as Python, R, or Matlab. Video acquisition to save files was implemented with the PiCamera Python package and video streaming was achieved using the UV4L software. Video files are initially saved in the h264 format and automatically (and optionally) converted to the mp4 video file format.

### Swarm Software

The Swarm software is designed to provide a web interface for the control of all aspects of any number of behavior boxes equipped with a Raspberry Pi running the PiE server software. All functionality of one behavior box with a PiE server is included in the Swarm software. The Swarm software can be run on any remote (analysis) computer including Raspberry Pi, macOS, and Windows operating systems. The Swarm software connects to individual PiE servers over the internet and itself provides an easy-to-use web interface.

The Swarm software also allows automatic synchronization of recorded video files, copying them to a remote analysis machine for further analysis and safe storage. In addition to full experimental configuration and control, the Swarm software allows experimenters to review recorded files and view real-time live video streams. With this design, fully remote control and review is possible, thus reducing potentially confounding experimenter interaction with animals and allowing experiments to be controlled from remote locations such as other labs, while at home, or even abroad.

Once installed on a local or remote computer, the Swarm software runs a web-server (Flask) and provides a point-and-click GUI in a browser (JavaScript). All communication with individual PiE servers is through their built-in RESTful API.

### Video Annotation Desktop Software

We have implemented desktop software, called VideoAnnotate, to perform behavioral analysis. Importantly, VideoAnnotate will work with any video, it is not dependent on video acquired with the PiE server software. At its core, VideoAnnotate provides video playback including play, pause, playback rate, and random access to any time within the file. As video is reviewed, the keyboard is used to indicate the start and stop of any number of different behavioral events. Point behavior event markers are also provided. All events within a video are presented in an intuitive GUI for review and curation. For example, clicking on an event will play the video corresponding to just that event.

A key design feature of the VideoAnnotate software is that it includes a randomization algorithm to reduce experimenter bias. This also allows random sampling of longer videos to reduce analysis time while not losing statistical power. In addition, this randomization is important because behavior often depends on the sequence of events such as ‘just placed in a cage’. By presenting the video in a random sequence of Chunks (see below) to the analyzer, we reduce bias as the analyzer does not know the sequence of events they are viewing. Once analyzed, all VideoAnnotate events are saved as a comma-separated-value (CSV) text file that can easily be opened and further analyzed with custom analysis scripts in almost any language.

As an example, we divided video files for our behavioral experiments into three Pieces of 10 minutes each, then 10 Chunks of video (10 seconds/Chunk) from each Piece were randomly selected, to total 30 Chunks of behavior video analyzed per 30-minute recording. With this system, a 30-minute behavioral video can be analyzed blind, while only having to score 5 minutes of video. These parameters are all user-configurable in the VideoAnnotate desktop application.

### Maternal Behavior Tests

Lactation day 0 (LD0) was demarcated as the first day live pups were seen in the cage by 12 PM. Behavior experiments were conducted on Lactation Days 1, 3, and 5 (LD1, LD3, LD5) during the first hour of the light cycle from 09:00-10:00 AM. Immediately prior to the Maternal Behavior Test, the mouse dam was removed from the cage and placed in a clean separation chamber for 15 minutes. The cage was then cleaned, and food and water removed for the assay. Just before the start of the assay, the pups were returned to the cage and placed in the three corners opposite a new nestlet. The video recording was begun at the dam’s moment of return to the cage, which commenced a 30-minute video capture period.

### Behavioral Scoring

The 30-minute Maternal Behavior Test video was scored blind, consistent with standard maternal behavior testing (Numan and Callahan, 1980; Matthews-Felton *et al*., 1995; Felton *et al*., 1998; Nephew and Bridges, 2011; Wu et al., 2014). Nine parameters were scored as either present or absent in each 10-second randomly presented video Chunks as detailed in the video annotation software section. These were: all pups retrieved and grouped, all pups nested, pup retrieval, pup interaction (sniffing, anogenital licking), crouching over pups, resting with pups, nest building, solo activity, solo rest. Three additional parameters were scored by viewing the entire 30-minute Maternal Behavior Test as a whole: delay to initiate first pup retrieval, delay to complete pup retrieval, and nest score at 30 minutes. Nests were scored 0 to 4 as follows: 0, no nest attempted; 1, poor nest, not all of the nesting material was used, lacks structure; 2: fair nest, all the nesting material was used but the nest lacks structured walls; 3, good nest, all nesting material was used and the nest has low walls; 4, excellent nest, all the nesting material was used and the nest has high structured walls (Numan and Callahan, 1980).

### Statistical Analysis

All behavior data was analyzed using the Wilcoxon Rank Sum Test, a non-parametric test of distribution differences in continuous data between two groups.

## Results

The PiE system allows for the monitoring and recording of mouse behavior while housed inside a home cage. Each home cage is enclosed within a behavior box. The individual behavior box design enables individualized light:dark cycle control, isolates behaving mice from neighboring cages, and reduces mouse exposure to the experimenters. Early versions of the PiE system were successfully used to monitor home cage wheel running (Cudmore *et al*., 2017) and activity levels during the light:dark cycle (unpublished observation).

The PiE system is modular, starting with an individual behavior box, its sensors and actuators, and a dedicated Raspberry Pi computer running the PiE server software (**Figure 1a**). Because the PiE server software is running both a web- and file-server, complete remote control of the box can be achieved using the provided web interface from any web browser. This web interface allows remote-control of sensors, actuators, and video recording as well as live-streaming video and downloading recorded video files.

**Figure 1.**
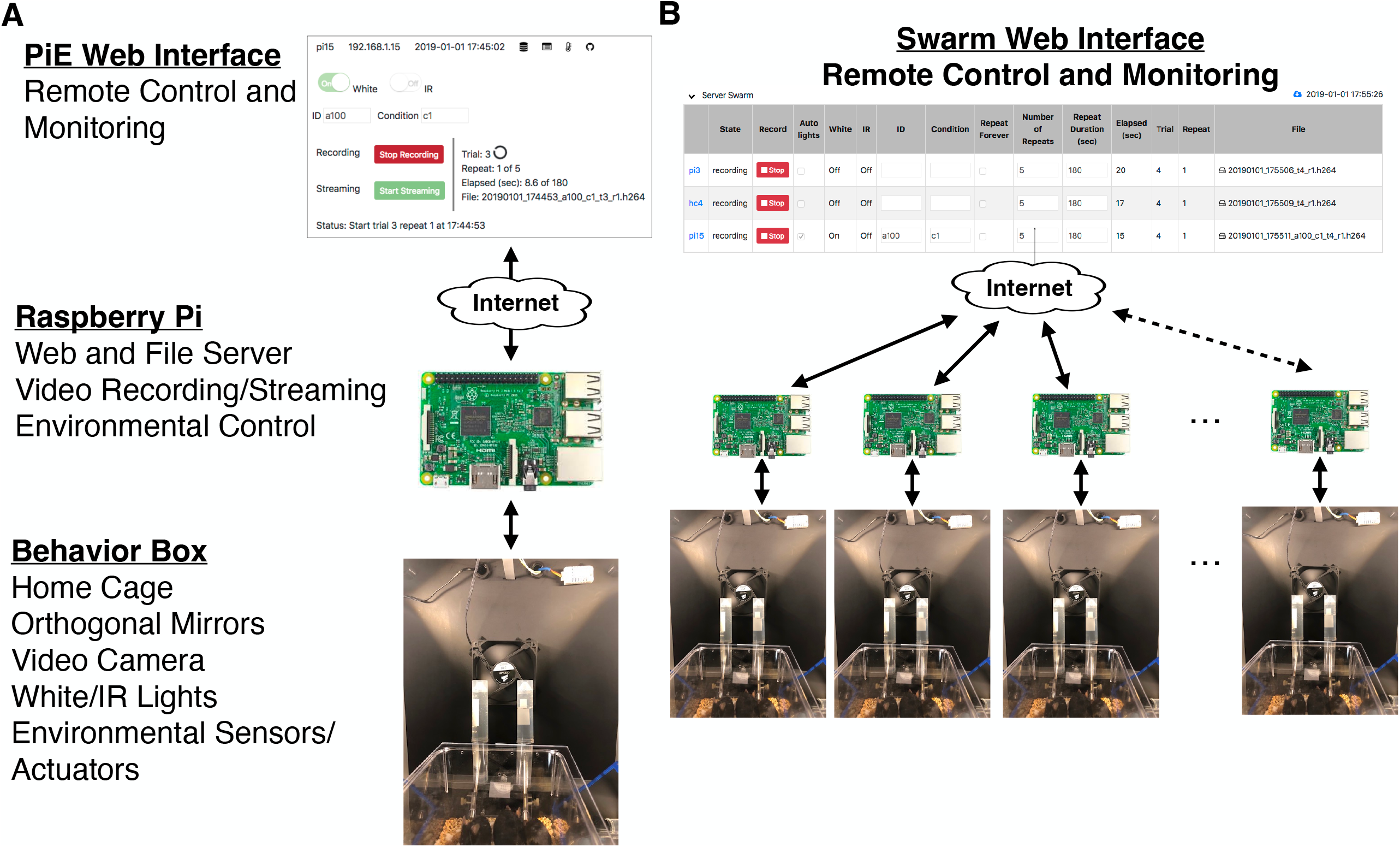
Overview of the PiE system. **(A)** Schematic of system for one behavior box. Each behavior box contains a home cage, video camera, day/night lights, and configurable environmental sensors and actuators (Behavior Box). Each box has a dedicated Raspberry Pi computer running the PiE server software which provides a web and file server, video recording/streaming, and environmental control of sensors and actuators (Raspberry Pi). The entire system can be configured, controlled, and monitored remotely via a web browser (PiE Web Interface). **(B)** Any number of PiE server equipped behavior boxes can be remotely controlled and monitored from the Swarm web interface. This includes setting all experimental parameters of each PiE server as well as monitoring a live video feed and transferring video from the PiE server to a remote analysis computer.

To scale up a behavioral experiment, one needs to include any number of behavioral boxes such that independent measurements can be made simultaneously over a large number of individuals, or in our case, mouse dams (mothers) with their pups. To achieve this, we implemented the Swarm web interface to remotely control any number of boxes running the PiE server software (**Figure 1B**). The Swarm web interface provides the same control of an individual PiE server interface just at the scale of an arbitrary number of behavior boxes.

We constructed light-proof boxes with off the shelf components designed to enclose a home cage (**Figure 2A**,**B**). Each box was internally equipped with sensors and actuators including a video camera, temperature and humidity sensors, cooling fans, and daytime (white) as well as nighttime (IR) LEDs (**Figure 2C**). To control the sensors and actuators within each box, a Raspberry Pi computer was attached to the back (**Figure 2D**). Video recording is achieved with a downward facing camera providing an aerial overview of a home cage (**Figure 2E**). In addition, we placed mirrors next to the home cage to provide left and right side-angle (orthogonal) views of the cage to improve behavioral scoring. By design, boxes can be stacked to allow high throughput behavioral analysis of multiple home cages simultaneously (**Figure 2F**).

**Figure 2.**
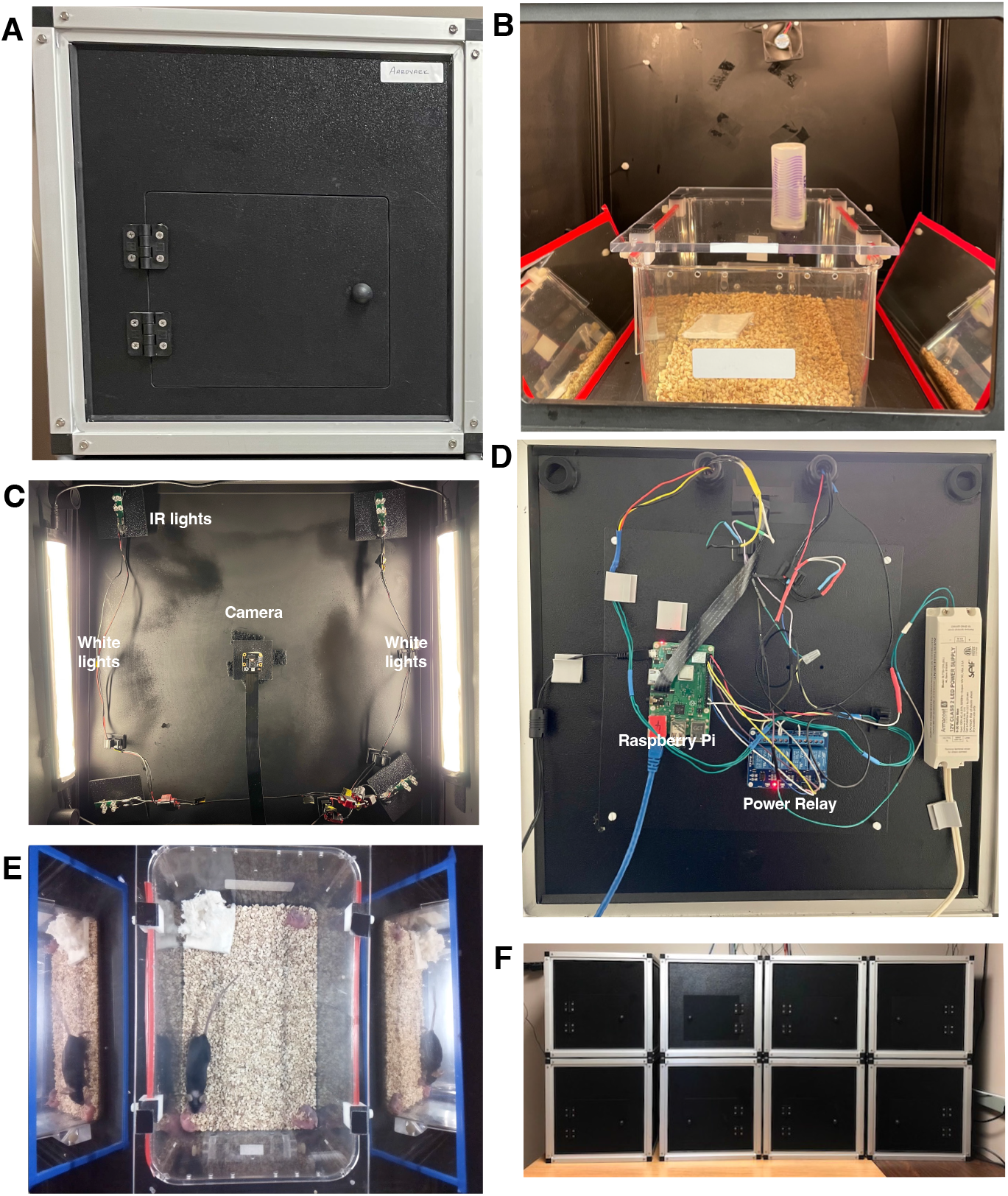
PiE behavior boxes. **(A)** Photo of the front of a behavior box including the door to access the inside. **(B)** Detailed picture inside one behavior box showing a home cage with a little of mice and a clear acrylic lid (lower), custom water dispensers for a sucrose preference assay (rear). This box also contains a temperature/humidity sensor (white rectangle in upper right), a cooling fan (center), white and IR lights, and a video camera (out of view). **(C)** View inside the box looking up at the celling. The camera is in the center of the image, white (daytime) lights are on the left and right and in each corner are IR LEDs for nighttime video. **(D)** Example wiring of a Raspberry Pi computer (green rectangle) and a relay to provide power to the lights (blue rectangle). This is attached to the back of each behavior box. **(E)** Example frame of acquired video with mom, pups, and bedding. Also shown are two mirrors at a 45-degree angle providing an orthogonal side-view of the cage to improve behavior analysis. **(F)** An example configuration of 8 PiE behavior boxes.

To ensure stable temperature and humidity within the enclosed (light tight) behavior boxes, we implemented a system of cooling fans and lightproof air vents. By continuously measuring temperature and humidity (using the PiE server software), we were able to ensure a stable environment.

The PiE system is a software pipeline that takes a behavioral experiment from raw data collection, to data organization and storage, to final analysis and plotting of the results. Here we describe each piece of this modular design including (i) PiE server software, (ii) Swarm software, and (iii) video annotation software.

### PiE server software

This software runs on an individual Raspberry Pi computer and provides both a web- and file-server. Throughout the development of this system, we performed extensive testing and modifications to ensure it performs reliably and that it captures the general needs of a remotely controlled behavioral experiment. The provided web interface allows all aspects of the behavioral data acquisition to be remotely controlled (**Figure 3A**). This includes starting and stopping of video recording as well as live streaming of the video feed. This also includes manual control of the daytime (white) and nighttime (IR) LEDs as well as the fans installed in the box for environmental control. To allow semi-autonomous data acquisition, the web interface provides control of the parameters for video acquisition and environmental control (**Figure 3B**). This includes parameters for video acquisition such as video recording duration and the number of repeats, as well as video acquisition resolution and frame-rate. In addition, environmental variables can be configured in this web interface including the timing of light/dark period transitions.

**Figure 3.**
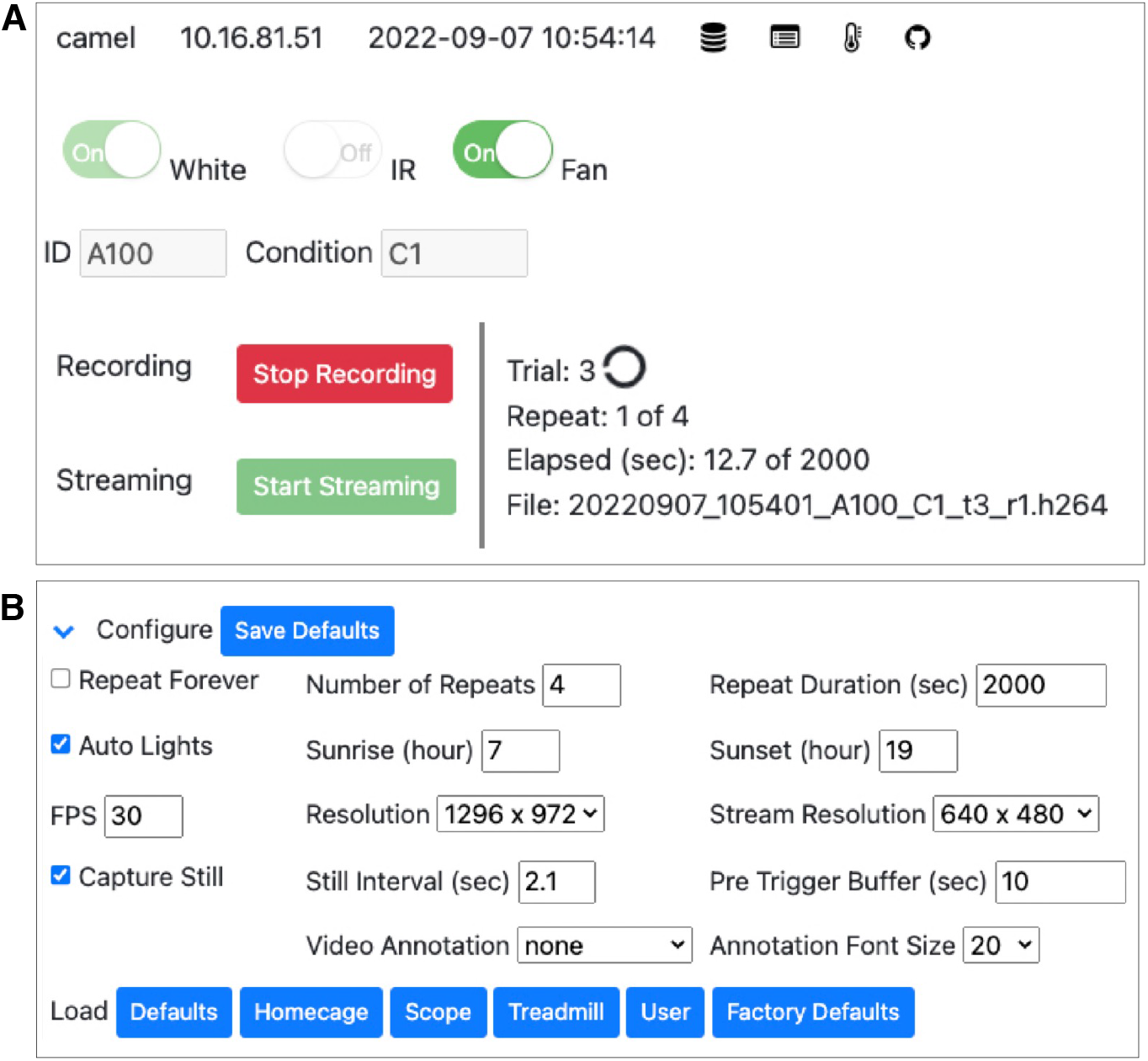
PiE server web interface. **(A)** Screenshot of the PiE server real-time web interface. This includes the name and IP address of the PiE server, its current date and time, and additional buttons to toggle different views including saved files, experimental logs, and environmental plots (top row). Controls are provided to manually toggle actuators such as white and IR lights as well as a cooling fan (second row). Experimental conditions such as animal ID and condition encodings can be entered (third row). Video recording and live streaming to the browser can be controlled (red and green buttons). The current status of the experiment is also shown including trial and repeat number, the elapsed recording time, and the current file being saved. **(B)** Each PiE server can be fully configured using the web interface. This includes video recording parameters such as the number of repeats and the duration of each recording, the timing of light/dark cycles, and video acquisition parameters such as frames-per-second and resolution. All options can be saved, and a number of option presets are provided (lower row, blue buttons).

As video and sensor data is acquired, it is saved locally on the dedicated Raspberry Pi computer attached to each behavior box. The PiE server software provides additional web interfaces to monitor this information and download it to a remote analysis computer. This includes a web interface to download recorded video files and textual logs of sensor data such as temperature and humidity (**Figure 4A**). Finally, the web interface provides plots and tabular logs of sensor data such as temperature and humidity (**Figure 4B**). This is critically important to monitor a behavior box remotely and can be used during offline behavioral analysis to determine if a disruption in these parameters occurred.

**Figure 4.**
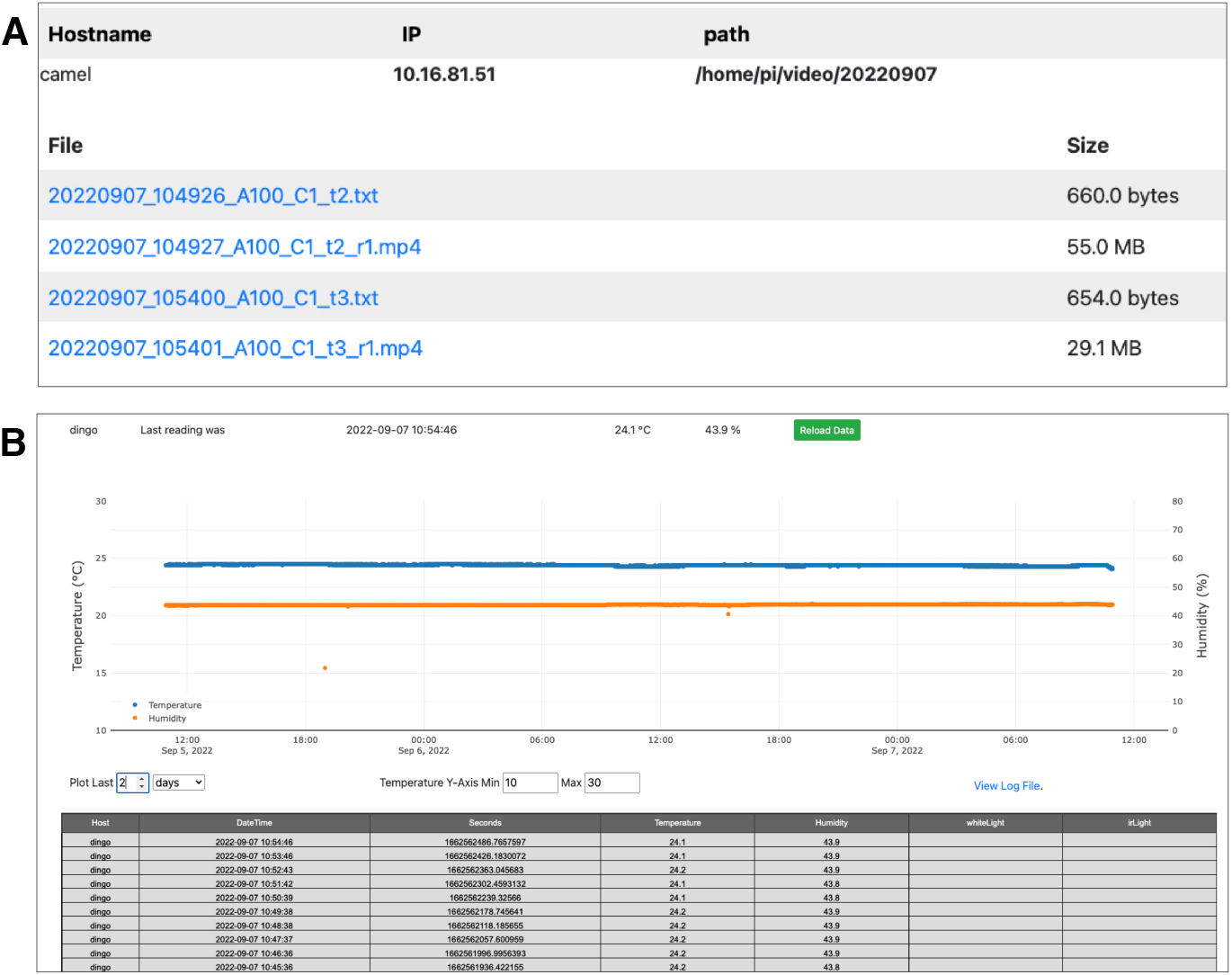
Additional PiE server web interface. **(A)** Each PiE server provides a web interface to review recorded video files and environmental logs. This interface allows individual files to be remotely viewed in a browser or downloaded to a local analysis machine. This example shows onetwo environmental log (txt) and fourtwo recorded video files (mp4). **(B)** A web interface is also provided to plot sensor data over arbitrary time-intervals. This example shows temperature (blue) and humidity (orange) over a one-month period. A table of all the raw data values is also provided (lower table). These values can be copied and pasted into a spreadsheet.

The PiE server software implements a web-based API to provide access to all its functions. This is called a RESTful API and basically provides a set of URLs to get and set all data and features of the provided web GUI. This includes an API to, for example, start and stop video recording, control actuators such as LEDs and fans, and to download acquired data. With this API, others can extend the capabilities of the PiE server software by creating customized scripts to implement novel behavioral data acquisition and control of the sensors and actuators included in the behavior box.

### Swarm software

Once an individual behavior box is configured and controlled with the PiE server software, a key feature of our pipeline is to extend this functionality to any number of behavioral boxes. This is achieved with the Swarm software that allows full control of any number of PiE servers to be remotely controlled (**Figure 5A**). The Swarm software provides the same control of individual PiE servers across any number of behavioral boxes. It allows for the video recording to be started and stopped, and for daytime and nighttime lights to be set to auto or manual. The Swarm software also provides a surveillance interface where the video recording and live feed of each behavior box can be monitored (**Figure 5B**).

**Figure 5.**
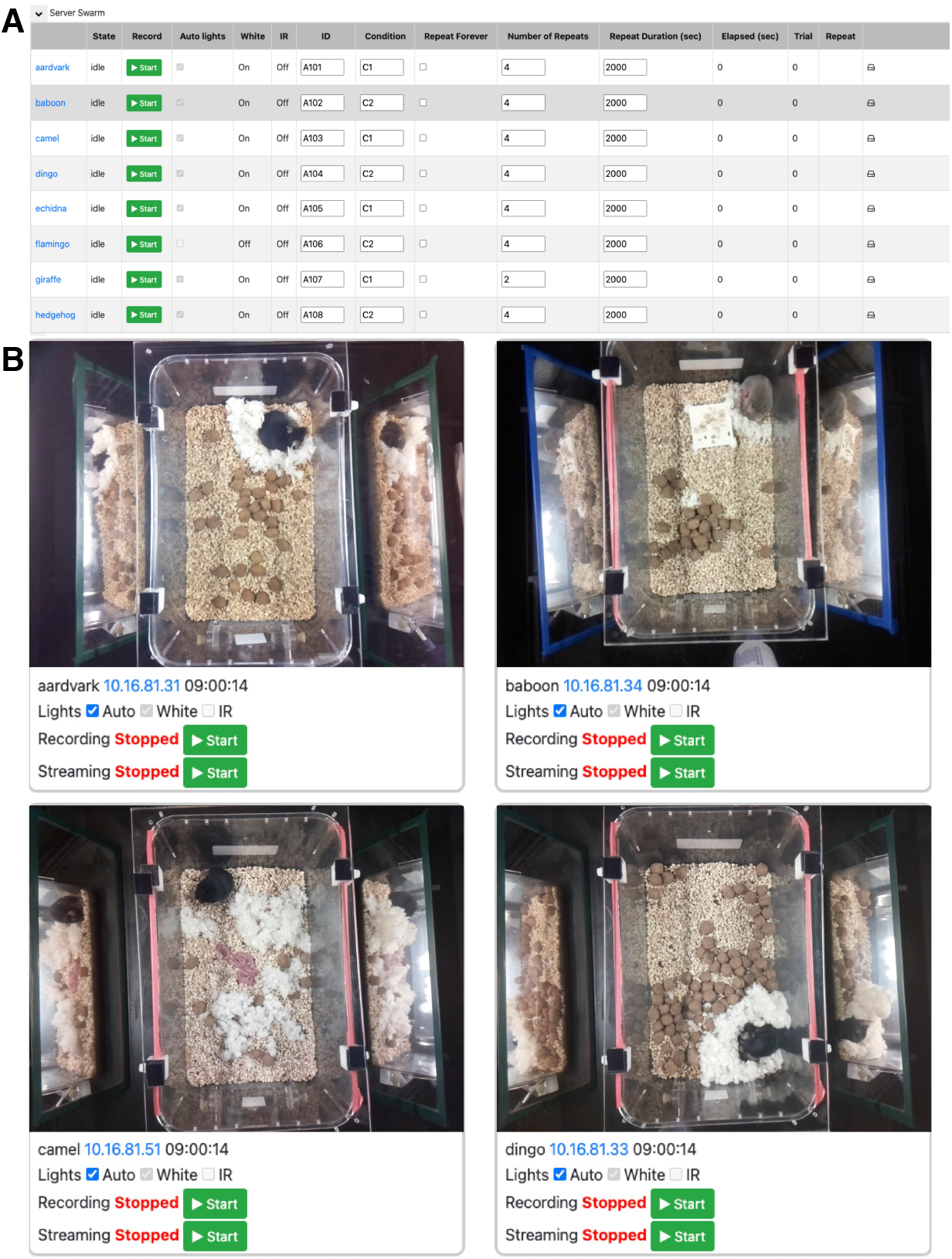
Swarm web interface to control any number of PiE behavior boxes. (**A**) An example screenshot of the Swarm web interface to remotely control an experiment and set parameters across any number of PiE server behavior boxes. In this example, eight PiE behavior boxes are being controlled (one per row). **(B)** A screenshot of the web interface to monitor and control any number of PiE behavior boxes with video recording and real-time video streaming. In this example, four PiE behavior boxes are being simultaneously monitored.

A key feature of the Swarm software is its ability to copy all raw data from any number of PiE servers to a remote analysis machine. This can be done while other experiments are running on the PiE servers and has an alarm-clock feature such that files can be automatically transferred at a given time or times per day. This is a requirement of our behavioral pipeline as just using 8 behavior boxes, we found manually collecting all the raw video data was tedious and error prone.

### Video annotation software

For the analysis of the raw video recorded and organized by the PiE server and Swarm software, we provide a desktop GUI for video annotation and scoring (**Figure 6A**). Unlike the PiE server software, this is designed to run on an analysis computer such as a macOS or Microsoft Windows operating system. At its core, this software interface provides all the expected video player capabilities including playing and pausing a video and random access to any frame of the video.

**Figure 6.**
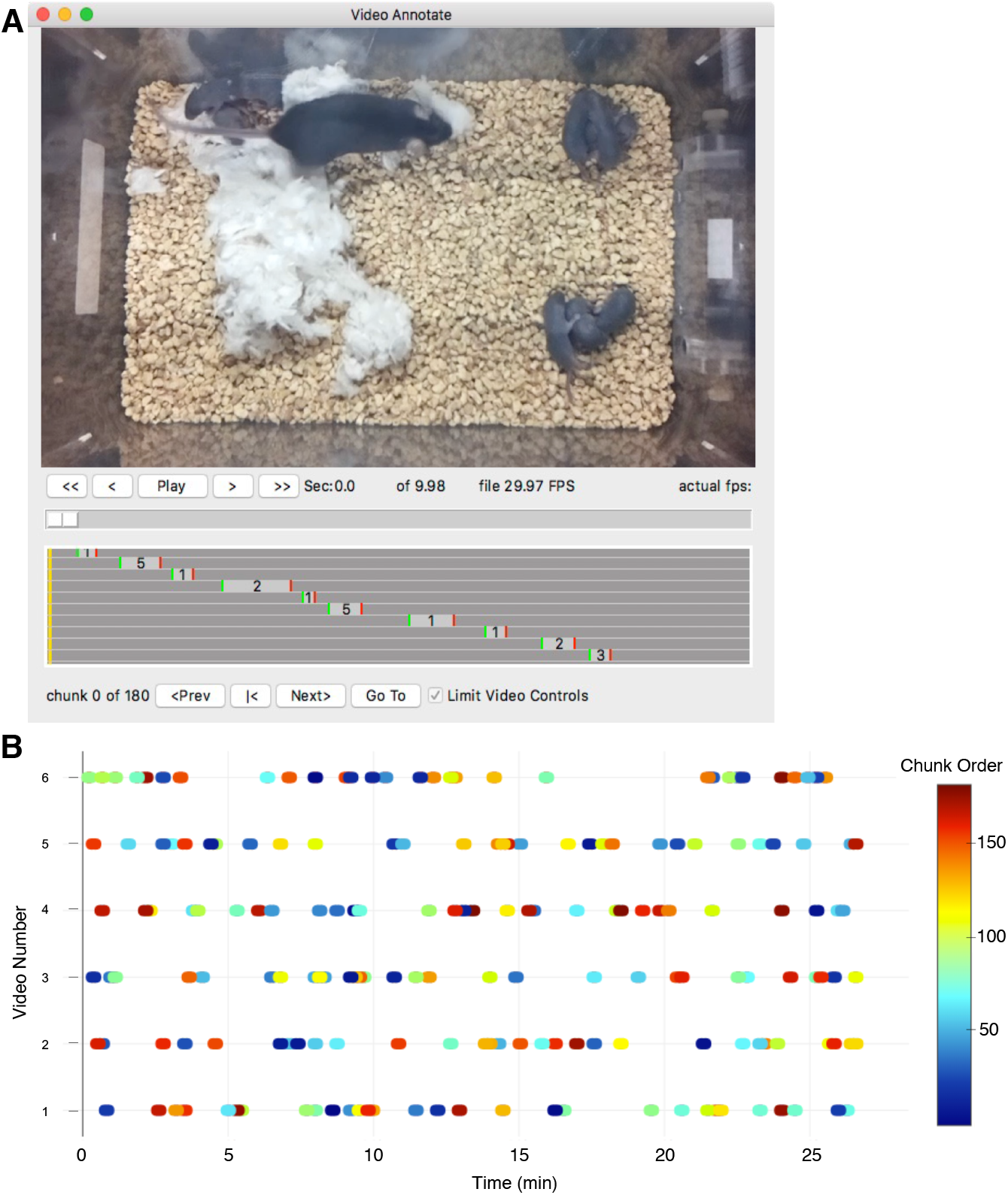
Video annotation desktop application. **(A)** Screenshot of the video annotation desktop application, VideoAnnotate. A video viewer is provided with controls to start, stop, and track within each video file (top). As behavior within the video is scored, the start, duration and type for each annotated behavior is dynamically displayed (gray box plot). When blinding the analysis with randomization, random Chunks can be presented in the main video window (bottom row of controls). **(B)** Example plot for one experiment showing the pattern of video Chunks automatically assigned for blind scoring. Six videos are shown (one per row) with each oval-symbol representing the time and duration of individual randomized video Chunks that were analyzed (30 Chunks for each file, 10 second Chunk duration, 180 Chunks in the total analysis). Color indicates the randomized order the video Chunks were presented in. The randomized algorithm also uses a Pieces parameter to seed the initial selection of Chunks (not show). For this example, three Pieces were specified with each Piece being 10 min of video (0-10 min, 10-20 min, 20-30 min).

The video annotation software is designed for the manual scoring of events within a recording. As the video is played, keyboard shortcuts allow the user to mark the beginning and ending of different behavioral events. This interface also allows the user to mark single time point behavioral events. All events are tallied and displayed in a GUI resembling a video editor interface.

Because behavioral experiments haveanalysis has the potential for experimenter bias, the video annotation software includes a randomization algorithm **(Figure 6B)**. For example, if the user has a 30-minute recording, this recording can be split into a desired number of Pieces (for example, 3 Pieces with a Piece duration of 10 min). VideoAnnotate then randomly selects an equal number of Chunks from within each Piece (for example, 10 Chunks per Piece with a Chunk length of 10 seconds). This allows for a total of 30 Chunks, with 10 Chunks from the first, middle, and last 10-minutes of the 30-minute assay. The Piece functionality is important for assays such as Maternal Behavior Testing, where certain behaviors, like pup retrieval or having all pups nested, are not equally likely to occur at the beginning or end of an assay. The three Pieces prevent the random sampling from randomly introducing a bias by selecting many Chunks towards the beginning or end of the assay. Though for assays where the behaviors scored are equally likely to occur at any point in the assay, Piece number should be set to “1” to nullify the Piece function. The user then proceeds to scoring, where VideoAnnotate presents each Chunk (pooled across all animals and experimental days the user wishes to score together), in a random order. With this system, videos of 30-minute duration can be scored in as little as 5 minutes. Finally, experimenter bias is reduced because the scoring is done in a random order and the scorer is blind to the experimental day, individual animal, and experimental group, as well as to the unique context of each Chunk within its total recording.

All behavioral events are saved as CSV text files which facilitate additional analysis and plotting by end users in any number of scripting languages. Importantly, the video annotation software does not require video to be acquired with the PiE server, it is designed to open, display, and analyze several common video formats including mp4.

## Exemplar Experiments

### Comparing maternal behavior in inbred mouse strains

To demonstrate the flexibility of the PiE system, we collected and analyzed home cage data using maternal behavior tests, a complex social behavior requiring simultaneous scoring of many behaviors (for a review of mouse maternal behavior, see Weber and Olsson, 2008).

Consistent with other work examining innate differences in maternal behavior in the DBA/2J and C57BL/6 inbred mouse strains, our data show significant differences between mouse dams of the two strains across a number of behaviors assayed (**Figure 7**). These measures included the delay to initiate pup retrieval (**Figure 7A**), scored video Chunks in which all pups were gathered in the nest (**Figure 7B**), and the nest score at the end of the assay (**Figure 7C**). These tests illustrate the different uses of the video annotation software including the analysis of complete video files (**Figure 7A**), analysis of randomly selected Chunks of video (**Figure 7B)**, and analysis of the end of each assay (**Figure 7C**).

**Figure 7.**
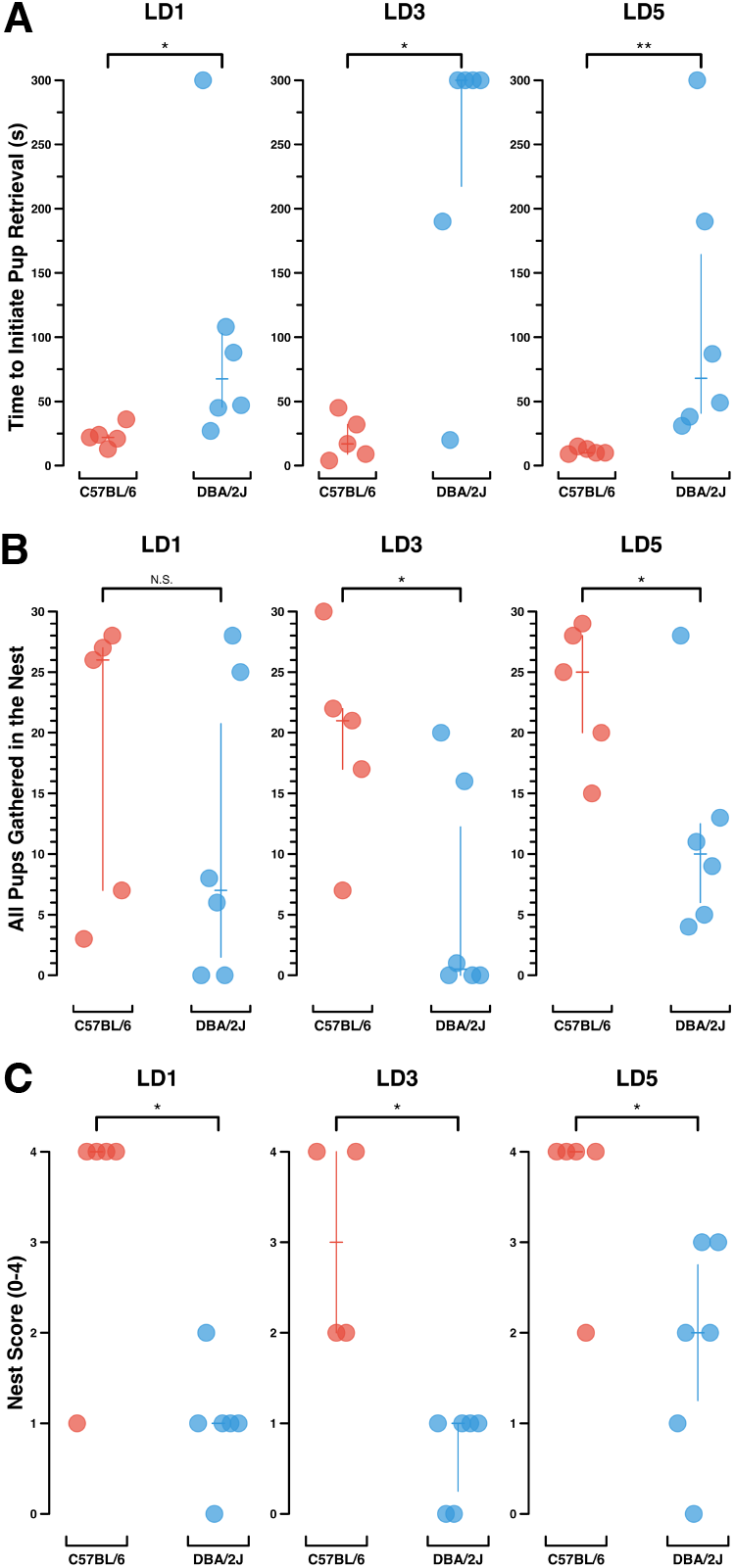
Maternal behavior analysis. Comparison of maternal behaviors in two inbred mouse strains, C57BL/6 (n = 5) and DBA/2J (n = 6). Each symbol (circle) represents a single dam’s maternal behavior performance on a given lactation day. Vertical bars depict the median ± interquartile range **(A)** Mouse dams’ time to initiate pup retrieval (s), up to a cutoff time of 300 seconds, on Lactation Days 1, 3 and 5 (LD1, LD3, LD5). Wilcoxon rank sum test, p-values for LD1, LD3, LD5: p = 0.016, p = 0.019, p = 0.008. **(B)** The number of video Chunks (out of 30 scored) in which the dam had all her pups gathered in the nest. Wilcoxon rank sum test, p-values for LD1, LD3, LD5: p = 0.36, p = 0.034, p = 0.044. **(C)** Nest score at the conclusion of a 30-minute maternal behavior assay, 0-4. Wilcoxon rank sum test, p-values for LD1, LD3, LD5: p = 0.024, p = 0.011, p = 0.03.

Across all three behavioral measures, DBA/2J dams show reduced maternal responsiveness as compared to C57BL/6 dams. DBA/2J dams take longer to initiate pup retrieval, they have all of their pups retrieved and nested in fewer of the video Chunks scored, and they build weaker nests than C57BL/6 dams. These findings are consistent with past findings comparing DBA/2J and C57BL/6 mouse strains (Carlier *et al*., 1982; Cohen-Salmon *et al*., 1985; Brown *et al*., 1999; Shoji and Kato, 2006).

## Discussion

We have designed an end-to-end pipeline to house animals in their home cage, allowing robust and reliable behavior, the ability to record 24/7 video, and analyze the resultant video in an unbiased way. A key feature of this system is that all aspects of an experiment can be controlled and monitored remotely with an easy-to-use web interface. This effectively reduces disruptions on behavior and will lead to more reliable and reproducible behavioral analysis.

This system is entirely open-source, all code and construction recipes are included here and expanded in our online resources. By implementing all code in widely used languages including Python, R, and JavaScript we are confident that this system can be extended by others. From a hardware perspective, our system is scalable and fully extendable by others. This is facilitated by the increasing availability of low-cost off the shelf components (e.g. Sparkfun, Adafruit, Digikey) but also components designed by other researchers and made available on public resources such as OpenBehavior and Hackster. By choosing the Raspberry Pi as the main hardware controller, this is also open-source and extendable as the not-for-profit Raspberry Pi foundation is dedicated to educating new and seasoned users in using their computers and associated software.

Behavioral analysis is often fraught with reproducibility errors. This is partially due to small sample sizes but also attributed to unintentional changes in an animal’s environment including interactions with the experimenter and neighboring researchers and moving animals to a novel and stressful environment. It is important that confounding variables be reduced, controlled, and eventually abolished. We have achieved this by keeping mice in their home cage in a dedicated environmentally controlled box that is remotely controlled through a web-browser, allowing the experimenter to be in another room, out of earshot.

Our behavioral analysis of recorded video is manual. While machine learning algorithms for scoring maternal behavior and other complex social behaviors are continuously improving, for many experiments the data scale and time required to train a machine learning algorithm is not efficient. Currently, the cutting-edge of applying Deep Lab Cut to maternal behavior (Winters *et al*., 2022) can only successfully identify pup retrieval events with 86.7% accuracy. Agreement between two blinded scorers using PiE’s manual scoring software to identify pup retrievals was 100%. Machine learning algorithms for complex social behaviors are complicated by multiple freely-moving mice engaging in interdependent behaviors such as pup retrieval (to the nest) and nest building. To achieve this complex behavioral analysis, our video annotation software, VideoAnnotate, provides a robust workflow to both manually annotate behavioral events and to review and curate these events later or with other researchers. VideoAnnotate is general purpose and we expect it to be adopted by others as it does not depend on any particulars of the PiE system and only requires a video recording. Finally, we are eager for the continued improvement of machine learning algorithms to further enable other forms of complex analysis with the videos acquired using the PiE behavior boxes.

We have used the PiE system to record and analyze maternal behavior, a complex social behavior. Due to the sensitivity of rodent dams to stress (Nephew and Bridges, 2011), maternal behavior is an excellent use-case for conducting behavior experiments in the animal’s home cage with minimal experimenter and environmental disruptions. In our analysis we reproduce expected differences in maternal behavior between the DBA/2J and C57BL/6 inbred mouse strains. Consistent with other reports, C57BL/6 mice perform faster and more complete pup retrieval, and build stronger nests (Carlier *et al*., 1982; Cohen-Salmon *et al*., 1985; Brown *et al*.,1999; Shoji and Kato, 2006).

Taken together, the PiE system is an end-to-end behavior pipeline from home cage animal housing, to data capture and management, to behavioral scoring and analysis. Its modular design allows others to extend any component including the design of behavioral boxes, environmental control, behavior box data acquisition, and behavioral analysis of recorded video. These features make this open-source, remotely controlled, and user-configurable rodent behavior pipeline a logical choice for a low cost, flexible, and easy to use behavioral analysis pipeline.

## Acknowledgments

We thank David J Linden (JHMI) for continued support of this project and thank Jackson Feeny and the Johns Hopkins Biostatistics Center for their consultation.

## References

Bains, R. S., Cater, H. L., Sillito, R. R., Chartsias, A., Sneddon, D., Concas, D., et al. (2016). Analysis of Individual Mouse Activity in Group Housed Animals of Different Inbred Strains using a Novel Automated Home Cage Analysis System. Front Behav Neurosci 10. Available at: https://www.frontiersin.org/articles/10.3389/fnbeh.2016.00106 [Accessed August 30, 2022].

Bains, R. S., Wells, S., Sillito, R. R., Armstrong, J. D., Cater, H. L., Banks, G., et al. (2018). Assessing mouse behaviour throughout the light/dark cycle using automated in-cage analysis tools. J Neurosci Methods 300, 37–47. doi: 10.1016/j.jneumeth.2017.04.014.

Balzani, E., Falappa, M., Balci, F., and Tucci, V. (2018). An approach to monitoring home-cage behavior in mice that facilitates data sharing. Nat Protoc 13, 1331–1347. doi: 10.1038/nprot.2018.031.

Berman, G. J. (2018). Measuring behavior across scales. BMC Biology 16, 23. doi: 10.1186/s12915-018-0494-7.

Brown, A. E. X., and de Bivort, B. (2018). Ethology as a physical science. Nature Phys 14, 653–657. doi: 10.1038/s41567-018-0093-0.

Brown, R. E., Mathieson, W. B., Stapleton, J., and Neumann, P. E. (1999). Maternal behavior in female C57BL/6J and DBA/2J inbred mice. Physiol Behav 67, 599–605. doi: 10.1016/s0031-9384(99)00109-2.

Carlier, M., Roubertoux, P., and Cohen-Salmon, Ch. (1982). Differences in patterns of pup care in Mus musculus domesticus I: Comparisons between eleven inbred strains. Behav Neural Biol 35, 205–210. doi: 10.1016/S0163-1047(82)91213-4.

Cohen-Salmon, C., Carlier, M., Roubertoux, P., Jouhaneau, J., Semal, C., and Paillette, M. (1985). Differences in patterns of pup care in mice V—Pup ultrasonic emissions and pup care behavior. Physiol Behav 35, 167–174. doi: 10.1016/0031-9384(85)90331-2.

Cudmore, R. H., Dougherty, S. E., and Linden, D. J. (2017). Cerebral vascular structure in the motor cortex of adult mice is stable and is not altered by voluntary exercise. J Cereb Blood Flow Metab 37, 3725–3743. doi: 10.1177/0271678X16682508.

Felton, T. M., Linton, L., Rosenblatt, J. S., and Morrell, J. I. (1998). Intact neurons of the lateral habenular nucleus are necessary for the nonhormonal, pup-mediated display of maternal behavior in sensitized virgin female rats. Behav Neurosci 112, 1458–1465. doi: 10.1037/0735-7044.112.6.1458.

Genewsky, A., Heinz, D. E., Kaplick, P. M., Kilonzo, K., and Wotjak, C. T. (2017). A simplified microwave-based motion detector for home cage activity monitoring in mice. J Biol Eng 11, 36. doi: 10.1186/s13036-017-0079-y.

Goulding, E. H., Schenk, A. K., Juneja, P., MacKay, A. W., Wade, J. M., and Tecott, L. H. (2008). A robust automated system elucidates mouse home cage behavioral structure. Proc Natl Acad Sci U S A 105, 20575–20582. doi: 10.1073/pnas.0809053106.

Krakauer, J. W., Ghazanfar, A. A., Gomez-Marin, A., MacIver, M. A., and Poeppel, D. (2017). Neuroscience Needs Behavior: Correcting a Reductionist Bias. Neuron 93, 480–490. doi: 10.1016/j.neuron.2016.12.041.

Lauer, J., Zhou, M., Ye, S., Menegas, W., Schneider, S., Nath, T., et al. (2022). Multi-animal pose estimation, identification and tracking with DeepLabCut. Nat Methods 19, 496–504. doi: 10.1038/s41592-022-01443-0.

Luxem, K., Sun, J. J., Bradley, S. P., Krishnan, K., Pereira, T. D., Yttri, E. A., et al. (2022). Open-Source Tools for Behavioral Video Analysis: Setup, Methods, and Development. doi: 10.48550/arXiv.2204.02842.

Mathis, A., Mamidanna, P., Cury, K. M., Abe, T., Murthy, V. N., Mathis, M. W., et al. (2018). DeepLabCut: markerless pose estimation of user-defined body parts with deep learning. Nat Neurosci 21, 1281–1289. doi: 10.1038/s41593-018-0209-y.

Matthews-Felton, T., Corodimas, K. P., Rosenblatt, J. S., and Morrell, J. I. (1995). Lateral habenula neurons are necessary for the hormonal onset of maternal behavior and for the display of postpartum estrus in naturally parturient female rats. Behav Neurosci 109, 1172–1188. doi: 10.1037/0735-7044.109.6.1172.

Nath, T., Mathis, A., Chen, A. C., Patel, A., Bethge, M., and Mathis, M. W. (2019). Using DeepLabCut for 3D markerless pose estimation across species and behaviors. Nat Protoc 14, 2152–2176. doi: 10.1038/s41596-019-0176-0.

Nephew, B. C., and Bridges, R. S. (2011). Effects of chronic social stress during lactation on maternal behavior and growth in rats. Stress 14, 677–684. doi: 10.3109/10253890.2011.605487.

Numan, M., and Callahan, E. C. (1980). The connections of the medial preoptic region and maternal behavior in the rat. Physiol Behav 25, 653–665. doi: 10.1016/0031-9384(80)90367-4.

Redfern, W. S., Tse, K., Grant, C., Keerie, A., Simpson, D. J., Pedersen, J. C., et al. (2017). Automated recording of home cage activity and temperature of individual rats housed in social groups: The Rodent Big Brother project. PLOS ONE 12, e0181068. doi: 10.1371/journal.pone.0181068.

Richardson, C. A. (2015). The power of automated behavioural homecage technologies in characterizing disease progression in laboratory mice: A review. Appl Anim Behav Sci 163, 19–27. doi: 10.1016/j.applanim.2014.11.018.

Salem, G. H., Dennis, J. U., Krynitsky, J., Garmendia-Cedillos, M., Swaroop, K., Malley, J. D., et al. (2015). SCORHE: A novel and practical approach to video monitoring of laboratory mice housed in vivarium cage racks. Behav Res 47, 235–250. doi: 10.3758/s13428-014-0451-5.

Shoji, H., and Kato, K. (2006). Maternal behavior of primiparous females in inbred strains of mice: a detailed descriptive analysis. Physiol Behav 89, 320–328. doi: 10.1016/j.physbeh.2006.06.012.

Singh, S., Bermudez-Contreras, E., Nazari, M., Sutherland, R. J., and Mohajerani, M. H. (2019). Low-cost solution for rodent home-cage behaviour monitoring. PLOS ONE 14, e0220751. doi: 10.1371/journal.pone.0220751.

Spruijt, B. M., and DeVisser, L. (2006). Advanced behavioural screening: automated home cage ethology. Drug Discov Today Technol 3, 231–237. doi: 10.1016/j.ddtec.2006.06.010.

Voikar, V., and Gaburro, S. (2020). Three Pillars of Automated Home-Cage Phenotyping of Mice: Novel Findings, Refinement, and Reproducibility Based on Literature and Experience. Front Behav Neurosci 14. Available at: https://www.frontiersin.org/articles/10.3389/fnbeh.2020.575434.

Weber, E. M., and Olsson, I. A. S. (2008). Maternal behaviour in Mus musculus sp.: An ethological review. Appl Anim Behav Sci 114, 1–22. doi: 10.1016/j.applanim.2008.06.006.

Winters, C., Gorssen, W., Ossorio-Salazar, V. A., Nilsson, S., Golden, S., and D’Hooge, R. (2022). Automated procedure to assess pup retrieval in laboratory mice. Sci Rep 12, 1663. doi: 10.1038/s41598-022-05641-w.

Wu, Z., Autry, A. E., Bergan, J. F., Watabe-Uchida, M., and Dulac, C. G. (2014). Galanin neurons in the medial preoptic area govern parental behaviour. Nature 509, 325–330. doi: 10.1038/nature13307.

