## Supplemental Table 1 for "PiE: An open-source pipeline for home cage behavioral analysis"

### Materials and Equipment

| Product | Price at Publication | Notes | Link |
|---|---|---|---|
| Slotted Rails for Box Frames | Project dependent | | https://8020.net/20-2020.html |
| ABS Panels for Box Walls | Project dependent | | https://8020.net/65-2613-s.html |
| Transparent mouse cage bottom | Project dependent | | https://www.altdesign.com/mouse-cages/ |
| Acrylic for mouse cage tops | Project dependent | Optically transparent | https://8020.net/2602.html |
| Raspberry Pi 4 - 4GB RAM | 55.00 | This has a list of places to buy the latest Raspberry Pi | https://www.raspberrypi.org/products/raspberry-pi-4-model-b/?variant=raspberry-pi-4-model-b-4gb |
| MicroSD for Raspberry Pi OS | 9.95 | | https://a.co/3xWn2iS |
| USB for video storage | 14.69 | | https://a.co/d/a0OOtEt |
| Raspberry Pi NoIR Camera Board | 29.95 | V2 - 8 Megapixels available at time of publication | https://www.adafruit.com/product/3100 |
| Camera flex cable | 3.95 | This cable is preferable to the white one that comes with the camera because you want to minimize reflective surfaces inside the box | https://www.adafruit.com/product/2143 |
| 4-channel relay | 8.99 | | https://a.co/d/9MSxh6g |
| Jumper cables M-M | 1.95 | | https://www.adafruit.com/product/1956?gclid=Cj0KCQiAyoeCBhCTARIsAOfpKxgBxDelmI_MgZMxnHIIXkebpgL05hoSPHO7N2HIXqMe3hODA3IntfoaArIuEALw_wcB |
| Jumper Wires F-F | 3.95 | | https://www.adafruit.com/product/266 |
| Jumper Wires F-M | 3.95 | | https://www.adafruit.com/product/826 |
| Hook-up wire | 22.50 | Stranded wire is more flexible - solid core wire can be used here but we found stranded wire to be easier to solder. | https://www.sparkfun.com/products/11375 |
| Wire stripping tool | 7.99 | | https://a.co/d/13VGGXO |
| Soldering iron | 54.75 | | https://a.co/d/9kCbROy |
| Temperature Humidity Sensor | 7.95 | | https://www.adafruit.com/product/5181 |
| Power cords | 8.99 | For use from power source to Raspberry Pi and from power source to Quad Relay | https://a.co/d/ce8aGa2 |
| Wall power | 25.99 | For Raspberry Pis and Quad Relays | https://a.co/d/6WzYNTQ |
| Mirrors | 12.00 | | https://a.co/d/7w8RPo8 |
| Mirror adjustment stand | 4.95 | Test your configuration dimensions before building the exterior of the behavior box | https://www.adafruit.com/product/1679 |
| Fan | 12.66 | | https://www.digikey.com/en/products/detail/qualtek/FAD1-06025BBHW12-A/7724541 |
| Mounting Hardware | 14.95 | | https://www.adafruit.com/product/3658 |
| Ethernet Cables | 18.49 | | https://a.co/d/6SkX2uE |
| Router | 59.99 | | https://a.co/d/6ggbFR1 |
| Multi-port switch box | 43.99 | | https://a.co/d/9DOntxV |
| **Lighting:** | There are three different lighting options, outlined below: | | |
| **Option 1:** | Rigid LED bars | More expensive, Least Effort, Least Flexible | |
| White lights | 29.99 | | https://a.co/d/bJXB9DL |
| Infrared Lights | 15.95 | Note: These are 20” so you would need a box at least 21” wide for these to fit. | https://ledlightsworld.com/collections/ir-infrared-led-strips/products/12vdc-waterproof-ip65-smd3528-36-ir-infrared-850nm-940nm-led-linear-rigid-strip-36leds-3-6w-per-piece |
| **Option 2:** | LED Strips | Middle Expense, Middle Effort, Middle Flexibility | |
| White light strips | 13.99 | | https://a.co/d/8w5TzOP |
| Infrared LED strips | 35 | This single reel would cover multiple boxes so it’s more cost effective for multi-box projects than Option 1 but may be less cost-effective if the user is building a single box. | https://ledlightsworld.com/collections/ir-infrared-led-strips/products/dc12v-smd3528-300-ir-infrared-850nm-940nm-single-chip-flexible-led-strips-60leds-4-8w-per-meter?variant=17836968476762 |
| **Option 3:** | Individual LEDs | Lowest expense, most effort, most flexibility | Individually soldered |
| Printed circuit board | 2.99 | | https://www.superbrightleds.com/moreinfo/resistors/universal-9-led-pcb-sbl-pcb1/548/1743/ |
| White light LEDs | 6.95 | | https://www.adafruit.com/product/754?gclid=Cj0KCQiAyoeCBhCTARIsAOfpKxjF0gxa_VYT2nrAHeXsAAEzSJVIRxnQMbOP_eWktV09xGSgnAH3jzcaAhiWEALw_wcB |
| Infrared LEDs | 10.89 | | https://a.co/d/3pAPUmw |
| Splicers | 19.96 | User needs 4 per box, this is the price for 50 | https://a.co/d/7MjUhzW |
| Resistors | 0.95 | | https://www.sparkfun.com/products/14490 |
| Optional Upgrades: | | | |
| Mirror adjustment platform | 71.96 | Optional | https://www.thorlabs.com/newgrouppage9.cfm?objectgroup_id=10660 |
| Soldering workbench | 39.99 | Optional | https://a.co/d/b9er7TP |
| Camera tripod mount case | 2.95 | Optional | https://www.adafruit.com/product/3253 |
| Tripod mount | 9.84 | Optional | https://a.co/d/ihrs3sG |
| Forced air vents: | If active air exchange is preferred, an aquarium pump can be used to route airflow into the boxes through rubber grommets in the panels, with another tube and grommet on the opposite side of the box as an outflow vent | | |
| Aquarium pump | 14.99 | Optional | https://a.co/d/5mFpjFG |
| Rubber grommets | 6.99 | Optional | https://a.co/d/dOoeeqe |
| Black tubing | 23.78 | Optional | https://a.co/d/8YI0CPT |
